# Latent epistatic interaction model identifies loci associated with human working memory

**DOI:** 10.1101/016576

**Authors:** János Z Kelemen, Christian Vogler, Angela Heck, Dominique de Quervain, Andreas Papassotiropoulos, Niko Beerenwinkel

**Affiliations:** Department of Biosystems Science and Engineering, ETH Zurich, 4058 Basel, Switzerland; SIB Swiss Institute of Bioinformatics, 4058 Basel, Switzerland; Psychiatric University Clinics, University of Basel, 4012 Basel, Switzerland; Transfacultary Research Platform, University of Basel, 4055 Basel, Switzerland; Division of Molecular Neuroscience, Department of Psychology, University of Basel, 4055 Basel, Switzerland; Department Biozentrum, Life Sciences Training Facility, University of Basel, 4056 Basel, Switzerland; Division of Cognitive Neuroscience, Department of Psychology, University of Basel, 4055 Basel, Switzerland

## Abstract

**Background:** Epistatic interactions among genomic loci are expected to explain a large fraction of the heritability of complex diseases and phenotypic traits of living organisms.

Although epistasis detection methods are continually being developed, the current state of the art is exhaustive search methods, which become infeasible when the number of analyzed loci is large.

**Results:** We develop a novel latent interaction-based selection method for polymorphic loci as the first stage of a two-stage epistasis detection approach. Given a continuous phenotype and a single-nucleotide polymorphism (SNP), we rank the SNPs according to their interaction potential. When tested on simulated datasets and compared to standard marginal association and exhaustive search methods, our procedure significantly outperforms main-effect heuristics, especially in the presence of linkage disequilibrium (LD), which is explicitly accounted for in our model. Applied to real human genotype data, we prioritized several SNP pairs as candidates for epistatic interactions that influence human working memory performance, some of which are known to be connected to this phenotype.

**Conclusions:** The proposed method improves two-stage epistasis detection. Its linear runtime and increased statistical power contribute to reducing the computational complexity and to addressing some of the statistical challenges associated with the genome-wide search for epistatic loci.

## Background

With the advent of high-throughput genotyping technologies, huge bodies of genotype data have become available, holding the promise of deciphering the complexity of heritable traits and diseases. Genome-wide association studies (GWAS) have had remarkable success in unravelling the genetics of several Mendelian traits [1]. However, most phenotypes, including many diseases, have complex genetic causes, and it remains a challenge to explain their heritability [2]. Epistasis is thought to be the main genetic mechanism to connect the genotypic variability to phenotypes, as it encodes the molecular interactions between genes or gene products. With the growing size of current GWASs detecting genome-wide epistasis is a challenging task, posing computational and statistical problems alike.

The need for feasible epistasis detection tools has led in recent years to several methods being proposed for this task [3, 4]. The proposed epistasis search strategies generally fall into one of the following three categories: exhaustive, stochastic, or heuristic. Each type of method has its own advantages and disadvantages. Exhaustive methods are typically expected to perform most accurately, although correction for multiple testing often limits the statistical power. Indeed, comparison studies confirm their good performance, but also reveal a major drawback in runtime. For two-locus searches the exhaustive runtime is quadratic in the number of loci [5]. TEAM [6] and BOOST [7], two exhaustive search methods, have been found to exhibit the best epistasis detection power among five approaches compared, the remaining three being heuristic in nature [8].

The performance of stochastic methods is difficult to assess. They tend to find interesting results in GWAS data; yet, they often depend on main effects, as is the case for Random Forests [9]. Empirical comparison studies [10] have shown stochastic methods, such as Random Forests, to be outperformed by exhaustive search methods like Multifactor Dimensionality Reduction [11]. In addition, heuristic methods, such as, for example, Logistic Regression [12], often perform similarly or better than stochastic methods [10]. Similar conclusions have been drawn when comparing the stochastic EpiMode method [13] to BOOST and to the heuristic SNPRuler method [14], among others [3].

Epistasis detection can be regarded as a feature selection task, where the final goal is to extract complex multivariate features, namely interacting loci, from a model relating genotype to phenotype. In practice, to reduce the burden of high dimensionality, marginally associated or additive SNPs are often pre-selected in a first step. The heuristic methods using such greedy strategies, including SNPHarvester [15] or Screen and Clean [16], will inevitably discard those interacting SNPs that display weak main effects or depart from the additive model.

In the present study, we address the problem of bi-locus epistasis detection as a feature selection process. We introduce a novel method called Latent Interaction Modelling for Epistasis Detection (LIMEpi). While heuristic in nature, our approach departs from the additive or main effect-based (marginal) feature selection, accounting for potential nonlinear interactions. The method is formulated in the framework of structural equation modelling (SEM) [17, 18] and, in particular, moment structure analysis [19], using interactions with latent variables [20]. LIMEpi involves a latent variable regression for genotype-phenotype association and we derive analytical solutions for its parameter estimates. This achievement allows for ranking the SNPs in computation time linear in the number of SNPs, the same time complexity as for marginal association testing. With the analytical results at hand, we perform power analyses and assess the effect of the model parameters on the performance of the method.

We compare LIMEpi to the generic heuristic-marginal and exhaustive approaches, as reference points for the above described method classes, and present in detail the advantages and disadvantages across the parameter space. We find that the major advantage of our method resides in identifying epistatic SNPs with small or no main effects. As our analysis reveals, this situation can arise due to negative LD between the two causative SNPs. We investigate a continuous human working memory phenotype, as a case study, and use SNP array data to detect epistatic associations. LIMEpi, followed by an exhaustive search with the resulting candidate SNPs, prioritizes interactions between genes involved in the G-protein responses to neurotransmitters (*PDE1A*), neurodevelopment (*RBFOX1*), synaptosome (*SNAP-25*) and cell adhesion (*NRXN1*), among others, several with demonstrated function in working memory and related pathological conditions, such as schizophrenia.

## Results

### Statistical model

Although it is possible to accommodate more complicated epistasis models, we have chosen a simple instance of statistical epistasis [21] that allows for an analytical treatment. We consider the following relationship between the phenotype, *y*, an observed polymorphic locus, *x*, and a latent random variable, *z*, which we think of as an unknown SNP:

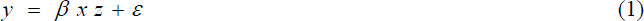

where *β* is the interaction coefficient and *ε* is an additive random perturbation of the phenotype. The model in Equation (1) is equivalent to the multiplicative model for case-control studies [22], and the assumed causal associations involved are summarized in the path diagram in Figure 1. LIMEpi estimates the parameter *β* (see Methods), which reflects the strength of the potential genetic interaction and tests whether the interaction is statistically significant. SNPs selected as capable of significant interactions are tested again in a second stage, this time against all genotyped SNPs, to identify their partners.

**Figure 1 -.**
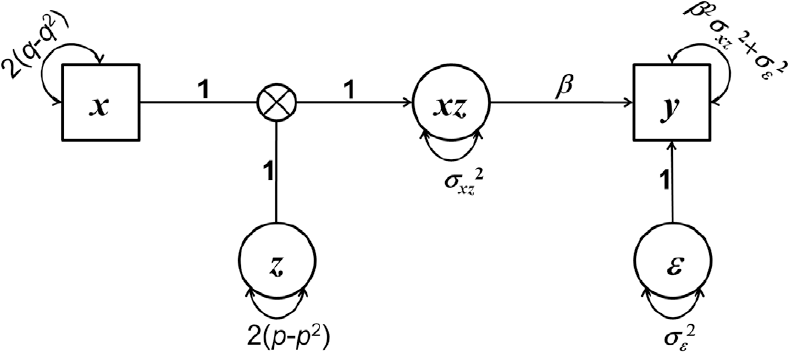
Path diagram of the model. Observed and latent variables in Equation 1 are depicted as squares and circles, respectively. Measured variables are the phenotype, *y* and the genotyped SNP, *x*. Latent variables are the latent SNP, *z*, the interaction variable, *xz* and the exogenous noise, *ε*. Single-headed arrows represent causal linear relationships and double-headed arrows represent variances. Parameters *q* and *p* represent the MAFs of *x* and *z*, respectively. 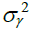 denotes the variance of the variable *γ*.

### Robustness to phenotypic noise

One factor that obstructs the detection of phenotype-associated SNPs or epistatic pairs of SNPs is the random perturbation on the phenotype. This noise can have measurement or environmental sources, such as other genetic factors or environmental conditions. In our model, all these sources are lumped into the random variable *ε*, which is assumed to be independent of both predictor SNPs, *x* and *z*. To assess the robustness to noise of LIMEpi compared to marginal detection and exhaustive search, we used the parametric *t*-statistic obtained from the analytical treatment of the three approaches as described in the Methods section, given the model in Equation (1). We express the *t*-statistic as a function of the exogenous noise, and compare it to the detection thresholds for the significance levels (nominal α = 0.05) Bonferroni corrected for 3,390 SNPs tested and for exhaustive pair-wise search, respectively (Figure 2A). The parameters we used for the analytical expression of the *t*-statistic were the following: the minor allele frequencies (MAFs) of the two causative SNPs were 0.2 and the LD between them was zero. The noise level is reported on the genetically meaningful scale of heritability in the broad sense [23], denoted *H*^2^, and computed as the variance of the interaction term divided by the variance of the phenotype. At a heritability of *H*^2^ = 0.02, the exhaustive approach performs above the significance threshold (Figure 2A). While both LIMEpi and the marginal association testing are below the significance threshold, LIMEpi has higher statistical power (Figure 2A). As the heritability increases to about 0.06, all three methods are able to detect the interacting SNPs.

**Figure 2 -.**
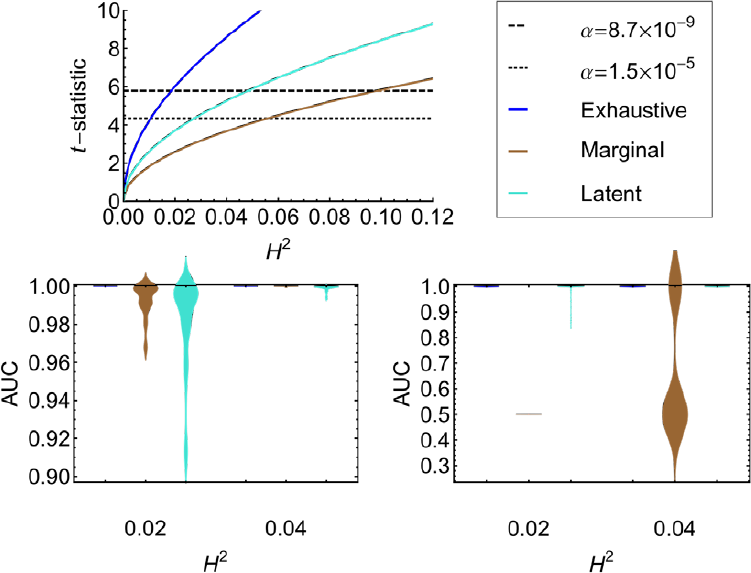
Performance as a function of heritability. Comparison of LIMEpi against the marginal and exhaustive methods at MAF=0.2, LD ≈ 0, 2,000 individuals, and 3,395 loci. **A**. Statistical power analysis through the *t*-statistic as a function of the heritability. Heritability increases with decreasing the exogenous noise variance. The Bonferroni corrected significance thresholds are shown for 3,395 loci and for 5,761,315 pairs of loci. **B**. Smoothed AUC histograms for the three methods in the first stage of the feature selection over 100 simulated experiments. The first-stage SNP ranking experiments are performed at two levels of heritability. **C**. Smoothed AUC histograms for the second stage of testing, given the first-stage features selected by the three methods under comparison in **B**.

We have controlled the family-wise error rate using Bonferroni correction in the analytical power analysis above; however, the actual SNP ranking performance may differ on actual data due to the overall LD structure and thus dependence of the statistical tests. To assess this performance, we analyzed the SNP rankings produced by the three methods on 100 simulated datasets. The phenotype was simulated using Equation (1), where the two SNPs were chosen from the datasets to closely match the above parametric settings (MAFs = 0.208 and LD = 0.0007). Gaussian noise was used to modulate the phenotype heritability. The AUC distributions from our ROC analysis confirm the expected best ranking performance of the exhaustive search (Figure 2B). When comparing LIMEpi to the marginal test, however, the latter shows higher ranking performance at both *H*^2^ = 0.02 and *H*^2^ = 0.04. To reconcile these seemingly contradicting results of our analysis, we also report the performance after running the second step of the epistasis detection (see Methods) using the candidate SNPs found significant in the first stage (nominal α = 0.05) and identifying their interaction partners in the dataset. The ranking of the SNPs found to actually interact with other SNPs (Figure 2C) in the second stage of testing confirms the predictions of the power analysis (Figure 2A). Here, LIMEpi has more statistical power. The main effect testing is unable to detect any SNPs at *H*^2^ = 0.02, while at *H*^2^= 0.04 it fails in more than half of the 100 experiments (Figure 2C). LIMEpi, on the other hand, did select the epistatic SNPs in the first stage; therefore, the second stage ranking is fairly good.

### Statistical power depends on the MAFs of the interacting SNPs

The variance of the interaction term in Equation (1) depends on the MAFs of the two interacting SNPs and so does, in turn, the heritability of the phenotype. We therefore expect a strong influence of the MAFs on the performance of epistasis detection.

We investigated the epistatic SNP detection power of LIMEpi versus the exhaustive and marginal approaches as a function of the MAFs (Figure 3A). The analysis was performed considering a model with two independent causative SNPs (LD = 0) with equal MAFs. The phenotype noise was set to the magnitude that yielded a heritability of 0.02 at MAFs of 0.2, as in the previous subsection. The statistical power of each of the three methods under comparison increases as the MAFs of the epistatic SNPs to be detected increase. LIMEpi has higher statistical power than the marginal detection over the entire range of MAFs, although at MAFs of 0.2 both methods are below the detection threshold (Figure 3A). For MAFs above 0.35, all three methods can detect interacting SNPs at the required significance level. Next, we analyzed the ranking performance of the three methods with respect to the MAFs on the 100 simulated datasets. To simulate the phenotypes we reused the two SNPs selected in the previous subsection (MAFs = 0.208 and LD = 0.0007) and then selected from the datasets two further SNPs with average MAFs of 0.5 and LD = 0.001. With MAFs of 0.5 the heritability of the phenotype increased to *H*^2^ = 0.1 at the exogenous noise level considered in the power analysis above. At this high level of heritability all three methods yield a perfect ranking of the interacting SNPs in the first stage of testing (Figure 3B). At MAFs of 0.2 however, we have the scenario of low heritability (*H*^2^ = 0.02), where the marginal test shows higher ranking performance in the first stage (Figure 3B). We performed again the second stage of epistasis detection using the SNPs called in the first stage (nominal α = 0.05) by the three methods. At a MAF of 0.2 the AUC distributions obtained with LIMEpi show significantly improved ranking performance over the main effect heuristic, which performs randomly at this MAF (Figure 3C). At a MAF of 0.5 all three methods rank the interacting SNPs accurately, as expected from the power analysis and the first stage of the procedure (Figure 3C).

**Figure 3 -.**
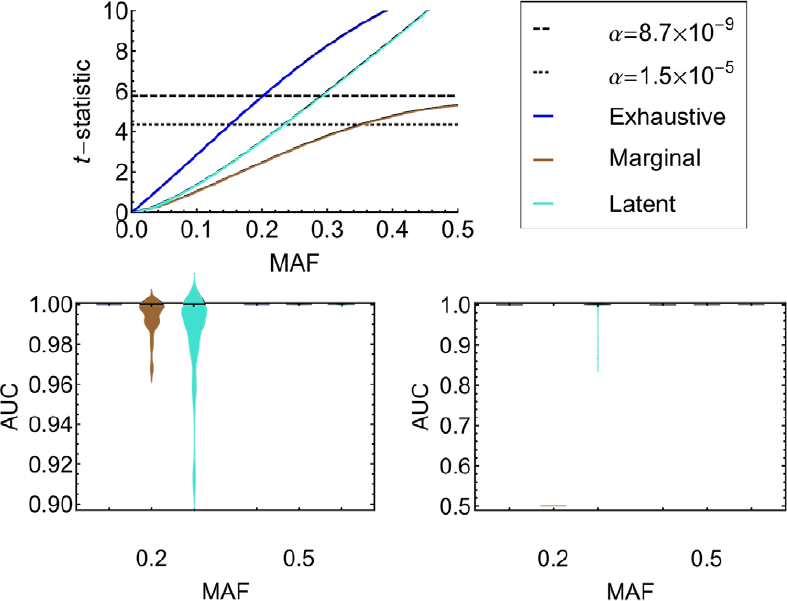
Performance as a function of the MAFs. Comparison of LIMEpi against the marginal and exhaustive methods at a noise level conferring heritability 0.02 at MAF=0.2, LD ≈ 0, 2,000 individuals, and 3,395 loci. **A**. Statistical power analysis through the *t*-statistic as a function of the MAFs of the causal SNPs. The statistical power increases with the heritability, which increases with the MAFs. The Bonferroni corrected significance thresholds are shown for 3,395 loci and for 5,761,315 pairs of loci. **B**. Smoothed AUC histograms for the three methods used as the first stage of the epistasis detection over 100 simulated experiments. The first-stage SNP ranking experiments are performed using two phenotypes simulated using two pairs of SNPs with MAFs 0.2 and 0.5, respectively. **C**. Smoothed AUC histograms for the second-stage of testing, given the first-stage features selected by the three methods under comparison in **B**.

### Strong negative LD implies pure epistasis

The impact of the LD between the causative SNPs on the statistical power to detect epistasis has long been recognized [22]. Here we analyze the effect of the entire range of LD on the performance of LIMEpi in comparison to the generic exhaustive and main effect methods. It is a strength of our modelling approach that it parametrically accounts for LD, even though we model and test only one genotyped SNP at a time.

We first investigated the expected effect of LD on the statistical power. The analysis was performed for a MAF of 0.5 to allow for the maximum range of LD (Figure 4A). The noise was kept at the same value as in the previous subsection, which yielded a heritability of *H*^2^ = 0.1 at LD = 0. Notably, the statistical power of LIMEpi is the same as that of the exhaustive search, while the marginal approach substantially drops in performance as LD approaches its lower bound (Figure 4A). To test whether the ranking performance is also consistent with this prediction, we applied LIMEpi to the simulated data, generating the phenotypes as follows. We searched the simulated datasets for SNPs with MAFs of approximately 0.5 and among those we selected the two SNPs with the largest negative correlation. The chosen pair had an LD of approximately −0.19. For the two selected SNPs, we simulated the phenotype using Equation (1). To assess the three methods’ performance for a different LD value, we reused the SNPs selected in the previous subsection (MAFs = 0.5 and LD = 0.001). We compared the AUC distributions for the three methods at the two levels of LD and observed excellent performance of LIMEpi both in terms of the true positive and false positive rate (Figure 4B). In contrast, the marginal detection method cannot cope with negative LD, since its underlying model does not account for LD in general. In practice however, zero or positive LD will allow marginal detection of interacting loci, as expected (Figure 4A) and observed on simulated data at LD = 0 (Figure 4B). The same conclusion can be drawn from the performance of the second stage of testing, where features selected in the first stage (nominal α = 0.05) are ranked in all possible pairs within the dataset (Figure 4C). Here, the main effect heuristic performs no better than random SNP selection. The lack of main effects due to negative LD strongly connects LD to pure epistasis, which cannot be detected through plain marginal nor additive effect detection.

**Figure 4 -.**
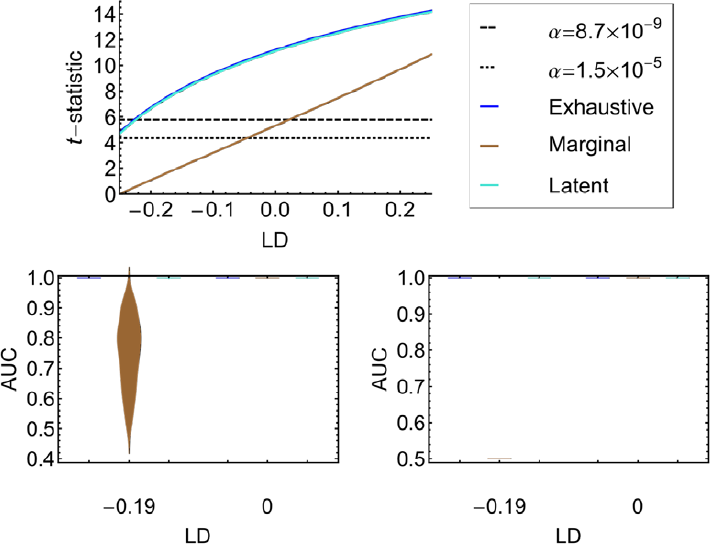
Performance as a function of LD. Comparison of LIMEpi against the marginal and exhaustive methods at MAFs 0.5 and a noise level conferring heritability 0.02 at MAFs 0.2 (LD = 0), 2,000 individuals, and 3,395 loci. **A**. Statistical power analysis through the *t*-statistic as a function of the LD between the causal SNPs. The statistical power increases with the LD. As the LD becomes negative, the marginal method will fail to pass the threshold. The curves for the exhaustive method and LIMEpi overlap. The Bonferroni corrected significance thresholds are shown for 3,395 loci and for 5,761,315 pairs of loci. **B**. Smoothed AUC histograms for the three methods used as the first stage of the epistasis detection over 100 simulated experiments. The first-stage SNP ranking experiments are performed using two phenotypes simulated using two pairs of SNPs at LD 0 and -0.19, respectively. **C**. Smoothed AUC histograms for the second stage of testing, given the first-stage features selected by the three methods under comparison in **B**.

### Runtime analysis

LIMEpi is based on a latent variable regression, which we solved analytically. For this reason, it has a linear time complexity in the number of loci, just like the marginal association testing procedure, which is performed here by linear regression of the phenotype on individual SNPs (see Methods). A major concern however, may be the computation time required for the second stage of testing. The second stage procedure chosen here exhausts all pairs-wise combinations between the SNPs from the selected set and all other SNPs. The second stage will therefore depend on the size of the feature set selected in the first stage. We performed an empirical analysis of the time complexity by investigating the number of SNPs selected by LIMEpi under various conditions.

First, we inspected how the overall LD in the datasets will influence the number of interaction candidates, since due to correlation we expect an increase of false positives. We used two samples of size 100 datasets each from the population of simulated genotype data taken at generations 500 and 750, respectively (see Methods). In generation 500, we observed a larger overall LD (average *r*^2^=0.034 compared to average *r*^2^=0.017 at generation 750) because the population size is much smaller at this time point. Since the exogenous noise, the MAFs of and the LD between the causal SNPs are parameters that influence the performance of LIMEpi, we simulated the several phenotypes to reflect the changes in these parameters. A two-fold increase of the overall LD produced an increase of the average number of selected SNPs by approximately two-fold in the worst-case scenario (MAFs = 0.5 and LD = 0.001) (Figure 5A).

**Figure 5 -.**
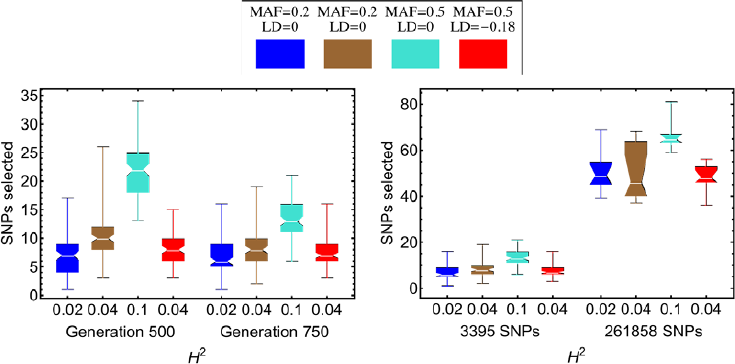
Computational complexity of the second stage. **A**. Distributions of the numbers of features LIMEpi identifies in the first stage at two overall LD levels within the simulated datasets. The overall LD decreases with the generation at which the sample was taken. To investigate how the complexity is affected by the parameters of the phenotype and causal SNPs (*H*^2^, MAFs and LD), we analyzed separately four scenarios where these parameters vary. **B**. Comparison between the numbers of features LIMEpi identifies in the first stage as the dimensionality of the simulated datasets increases. The complexity is inspected for different parameters as in **A**. The boxplots indicate the median, lower, and upper quartiles and the extreme values within one and a half inter-quartile ranges of the distributions of the number of selected SNPs.

Next, we investigated how the time complexity in the second stage scales with the dimensionality of the genotype dataset. For this purpose, we used a second whole-genome simulation of 500,000 loci (261,858 loci after quality control). We compared our feature selection method on ten datasets sampled from this population at generation 750 to the results obtained on the smaller scale data from the previous simulation at the same generation (3,395 loci after quality control). A 77-fold increase in dimensionality produced an average seven-fold increase of the second-stage computational complexity in the worst-case scenario (MAFs = 0.5 and LD = 0.001) (Figure 5B).

### Epistatic SNP candidate pairs involved in human working memory

Working memory, i.e., the ability to keep transitory information active and available for short-term manipulation and referencing during complex interactions with the environment, is a complex behavioural trait. Working memory performance is highly heritable and a number of genetic factors have been identified to contribute to its phenotypic variability [24]. These findings suggest new means of understanding and diagnosing closely related pathological conditions, such as schizophrenia [25]. A sample of 1947 young healthy Swiss individuals was assessed for working memory performance using an n-back task. All subjects were individually genotyped using the Affymetrix Human SNP array 6.0. The genotype dataset was filtered (MAF ≥ 0.1; HWE p_Fisher_ ≥ 0.01; call rate ≥ 95%) to obtain a set of 541,802 SNPs for further analysis.

We applied LIMEpi to the genotype dataset in a hypothesis-free fashion, using working memory as a continuous phenotype. We selected 132 candidate SNPs that passed the experiment-wide significance threshold (nominal α = 0.05, α_Bonferroni_ = 9.23·10^−8^). Subsequently, the second stage of testing was employed in order to identify actual interaction partners for these selected candidate SNPs within the dataset. We found 65 statistically significant pair-wise interactions (α_Bonferroni_ = 6.99·10^−10^). Out of these, 18 pairs had both interacting SNPs located in gene coding regions (Table S1). Among the 18 pairs are those involving *PLCH1*, a member of the eta family of the phosphoinositide-specific phospholipase C (PLC), and *PDE1A*, a Ca^2+^/Calmodulin-dependent phosphodiesterase (PDE). Both PLCs and PDEs are strongly implicated in the molecular mechanisms of working memory, such as the G-protein-coupled responses to neurotransmitters, which lead to Ca^2+^ release as the inositol trisphosphate concentration increases [26, 27], or cAMP/cGMP signaling in dopaminergic, cholinergic and serotonergic neurotransmission [28, 29]. Further significant interaction candidates include the zinc-dependent aminopeptidase *LNPEP* together with the xylosyltransferase *GXYLT1*, neurexin *NRXN1*, a cell adhesion molecule and receptor associated with schizophrenia [30] or *RBFOX1*, which regulates human neuronal development [31]. The SNP pairs for which only one locus resided in a gene coding region involved, among others, the synaptosomal-associated protein 25 (*SNAP-25*) and the proline rich membrane anchor 1 (*PRIMA1*), which anchors acetylcholinesterase at neural cell membranes.

The *p*-values reported in Table S1, show that none of the selected SNP interactions would have been found in an exhaustive interaction search with α_Bonferroni_ = 3.40·10^−13^. Among all the 65 statistically significant interactions, none were found significant with the Bonferroni correction corresponding to an exhaustive procedure. The smallest *p*-value observed was 1.14·10^−11^. We also performed a marginal association test between the phenotype and the individual SNPs. The smallest *p*-value observed was 2.43·10^−8^ and corresponded to SNP rs17628359 in *LNPEP* (encoding leucyl/cystinyl aminopeptidase), also identified by our proposed method. According to the marginal association test, no further SNPs were detected. The main effect-based heuristic would have been unable to declare further SNPs as significant, unless some arbitrarily higher significance threshold was used.

## Discussion

Latent variable models have been widely used to account for unobserved factors in biological systems and they have been shown to increase detection power in eQTL studies [32]. Here, we have introduced a novel latent interaction modelling approach, LIMEpi, to select candidate SNPs for epistasis analysis by assuming a latent SNP, which could modulate the association of an observed genotyped SNP with the phenotype. Through latent variable regression analysis we introduced and estimated a parameter, which accounts for the LD between the two potentially interacting genetic markers. Importantly, it has been reported in the population genetics literature that LD can be generated by epistasis and that multiplicative epistasis will maintain LD in a population [33]. Furthermore, recent studies indicate that LD can be used as a criterion for epistatic loci detection in case-control studies [34, 35]. Our method is applicable to both quantitative phenotypes, as exemplified in this paper, and to case-control designs using link functions [36]. Through careful analysis of the effect of LD on the statistical power to detect epistatic interactions, all in the context of our specific phenotype model and allele coding, we confirmed the tight connection between epistasis and LD. In fact, LD may be a hallmark of pure epistasis, since the total absence of marginal effects occurs only at extreme negative LD (Figure 4).

Our analysis has also shown the advantages and limitations of the method when the genetic heritability of the phenotype varies as a function of the phenotypic noise and of the MAFs of the genetic factors involved. The latent interaction approach was shown to have improved statistical power over the marginal association procedure. Moreover, when LD approaches its lower bound, the latent interaction scheme may even outperform state-of-the-art exhaustive search methods due to the more conservative Bonferroni correction required for the latter. The proposed first-stage SNP prioritization method scales linearly with the number of SNPs and, in practice, the runtime is comparable to that of marginal association detection, due to our analytical solution of the latent variable regression. The efficiency of this first stage of the two-stage testing will also lead to a significant time complexity reduction in the second stage of epistasis detection. In our genome-wide simulation study we noted an over 2,000-fold reduction in runtime compared to the exhaustive approach. For the real experimental data, we witnessed a similar reduction of the second-stage computational complexity.

We endeavoured to test the proposed method on an instance of real human data. The latent interaction modelling procedure (first stage) took around 20 minutes on a single 2.2 GHz processor core for a dataset of 1947 individuals and genotypic observations at 541,802 loci. The proposed first-stage feature selection is fast, suitable for large GWAS data, and hence contributes to solving the computational difficulties associated with genome-wide epistasis detection. We performed an exhaustive search with the candidate SNPs against the entire set of markers (second stage). At both stages we find reasonable amounts of statistically significant results, showing that we are able to overcome some of the statistical difficulties due to the high dimensionality of the data.

Specifically, we have found a likely interaction between a member of the PLC and a member of the PDE1 enzyme families. *PDE1A* has a higher affinity for the cGMP over cAMP. cGMP activates protein kinase G (PKG) which, in turn, has been shown to phosphorylate and regulate certain classes of G-protein-activated PLC (i.e., beta) [37]. The eta family of PLCs is the most recently discovered and studies suggest that at least some of its variants are G-protein-regulated [38]. Two other genes, namely *LNPEP* and *GXYLT1*, were also found to statistically interact. While none of them has been already associated with working memory, they both interact with HLA-B, the major histocompatibility complex, class I (MHCI), B protein [39-41]. Interestingly, MHCI has a role in synaptic remodelling and plasticity [42]. Since our study on the genetic background of the human working memory is ongoing, we expect future results with higher sample size to shed a clearer picture on the physiology and heritability of the complex cognitive trait of human working memory.

## Conclusions

We proposed a novel selection method for polymorphic loci, called LIMEpi, as the first stage of two-stage epistasis detection. We tested the method on simulated and real data and found that its linear runtime and improved statistical power contribute to reducing the computational complexity and to addressing some of the statistical challenges associated with the large-scale search for epistatic loci. We believe that our work will open up new possibilities in the analysis of epistatic interactions. These possibilities include, for example, the incorporation of additional latent interacting SNPs to target multiway interactions and extensions of the underlying model to more complex genotype-phenotype relationships [43, 44], although they may not allow analytical treatment anymore.

## Methods

### Latent variable regression

We denote the MAFs of the genotyped SNP *x* and the latent SNP *z* by *q* and *p*, respectively. The LD between *x* and *z* is defined as the deviation from random segregation of the alleles within the haplotypes [45], such that the expected haplotype frequencies are:

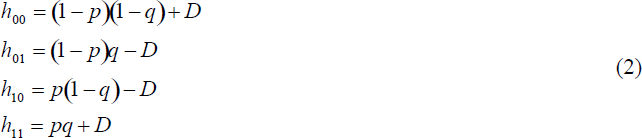

where *h*_*ab*_ denotes the frequency of haplotype *ab* (*a, b* ∈{0, 1}), where *a* is the allele status at the first locus (with genotype *x*) and *b* the allele status at the second locus(with genotype *z*). Assuming random mating of the two-locus haplotypes, we impose the Hardy-Weinberg Equilibrium (HWE) condition on the frequencies of the resulting two-locus genotypes. The joint probability of any two genotypes can be parameterized for all nine possible combinations so that the expected value of the product *xz* (*x, z* ∈ {0, 1, 2}) can be expressed parametrically in terms of *p, q* and *D*. *D* is a measure of LD, which also quantifies the covariance between the two individual SNPs with expected values E(*x*) = 2*q* and E(*z*) = 2*p*. From *h*_*ab*_ ≥ 0 in Equations (2) the following constraints apply to *D*:

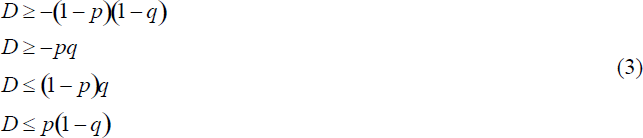

We analyze the model in Equation (1) as a structural equation model [19, 20] with three latent variables: *z*, *xz* and *ε* (Figure 1).

The four model parameters are *β, p, D*, and 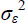, where 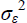 stands for the variance of *ε*. This structural equation model is non-standard, involving discrete and continuous distributions and an interaction effect. Inspired by [19], we use higher order moments as indicators of the latent interaction effect, to render the model identified. To estimate the model parameters, we apply constrained optimization to the weighted least-square cost function, as in an asymptotically distribution-free estimation [17, 46]. To shorten the runtime required by the numerical minimization procedure, we fixed *p* to 0.5, the value that confers maximum entropy to the latent variable *z*. Then the optimization problem becomes analytically tractable with parameters *β, D*, and 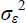.

There is no obvious optimal strategy to select the central moments for estimation. The main criteria are the identification of the model and the non-redundancy of the moments [17, 19]. Aiming for simplicity, we chose the following lowest order moments that rendered the model identified

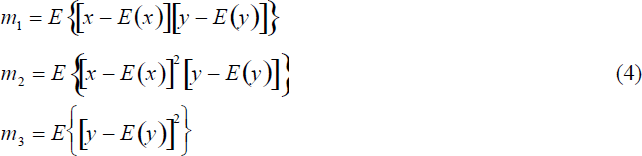

We use a saturated system of equations, such that the SEM problem can be regarded as a latent variable regression. The first moment, *m*_1_, is the covariance between *x* and *y*, and the third, *m*_3_, is the variance of *y*. These two moments are typically used in standard linear regression. Since we are using only the variance of *ε* here, we make no additional assumptions on the nature of the noise, except that it is independent of the predictor variables. Additionally, 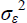 can be estimated from the expression for *m*_3_ in Equations (4) in a second step, once the parameters *β* and *D* have been estimated. Therefore, in the first step, we have a constrained system of equations involving only *β* and *D*. The constrains on *D* become box constrains, and the optimization problem to solve is

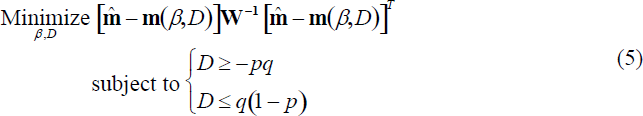

The weighted least-squares cost function in (5) involves the difference between the first two parametric moments in Equations (4) in the two-dimensional vector **m**(*β*,*D*) = (*m*_1_, *m*_2_) and their sample estimates denoted by the vector 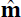 of the same dimension. The weight matrix **W** is the sample estimate of the asymptotic covariance matrix of 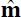 [17]. To solve the optimization problem (5), we used the Karush-Kuhn-Tucker optimality conditions [47] and solved the resulting system of quadratic equations analytically, using Wolfram Mathematica 8. The unconstrained solution is

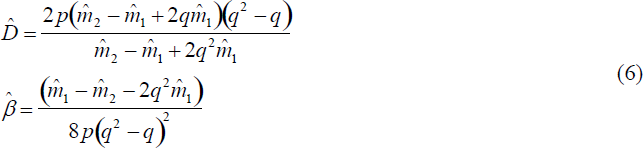

If the unconstrained solution is not feasible, two additional constrained solutions are evaluated at the boundaries of *D* and the one yielding the smallest cost solves the minimization problem (5). The analytical results were implemented in the R statistical software environment [48] and we developed the R package *LIMEpi*, to estimate the model parameters and to assess their statistical significance (see below).

### Marginal and exhaustive methods

The exhaustive analysis was performed in the form of a linear regressions using the same model as in Equation (1), but where both variables *x* and *z* are actual genotyped SNPs. For marginal association we used the linear model

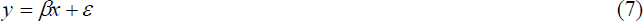

where *x* represents a single SNP.

### Statistical and power analysis

The statistical significance of an interaction or association was tested using the one-sample *t*-test of the null hypothesis that the interaction coefficient estimate is zero. To obtain standard error estimates for the parameter of interest we used White’s robust standard error [49],

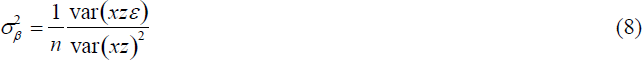

For the marginal regression models *xz* becomes the respective predictor variable, *x*. The sample size is denoted by *n*. The *t*-statistic and the degrees of freedom are then given by *t* = *β*/*σβ* and df = *n* − 1, respectively. For the power analysis, we took a model-based approach. We consider the true phenotype model to have the fixed parameters *β* = 1, *p, q, D* and 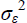. We assume that the exhaustive method will yield precise estimates 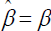 and 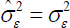 since both SNPs are available. For LIMEpi and the marginal approach we consider that only one SNP with MAF *q* is available. We express the moment estimates, 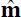 (Equations (4)) in terms of the true model parameters:

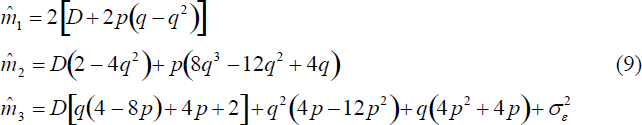

LIMEpi yields the estimates 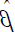 and 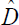 from Equations (6) and the corresponding 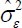. Linear regression on *x*, for the marginal approach, yields 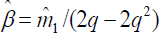 and the corresponding 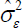. The estimates from each of the three methods are then used to compute the standard errors (Equation (8)) and *t*-statistics. We study the behaviour of the *t*-statistic as the parameters of the true model, *p, q, D* and 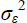 vary.

### Two-stage detection of epistatic SNP pairs

To identify epistatic SNP pairs with respect to a phenotype in a dataset of *L* genomic loci, we employ a modified Either Significant Two-Stage Strategy [51]. In summary, in the first stage, we apply our latent variable regression and obtain a set of *α*_1_ single SNPs with statistically significant potential to interact (nominal α = 0.05). In the second stage, we test each SNP selected in the first stage against all other loci within the dataset. To assess the statistical significance of the pair-wise association of SNPs with the phenotype, we compute the standard error of the regression coefficient estimates, as in Equation (8), and evaluate the *t*-statistic by computing the *p*-value. Because the second stage of testing is not independent of the first stage of feature selection, we compute conditional *p*-values for SNP pairs. We consider simultaneously both the latent interaction model and the interaction model with two true SNPs.

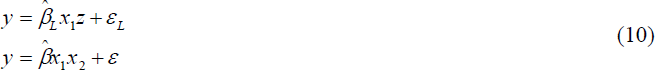

where *y* is the phenotype, *x*_1_ and *x*_2_ are two observed SNPs from the dataset, *z* is a latent SNP, 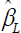 -and- 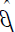 are estimates of the interaction strengths of the latent and the two-SNP interaction, respectively, and similarly, *ε*_*L*_ and *ε* are noise variables. We assume that the interaction coefficients *β_L_* and *β* follow a bivariate normal distribution:

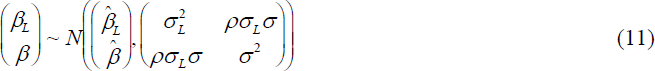

where *ρ* is the correlation between the two interaction coefficients and *σ*_*L*_ and *σ* represent their standard deviations, estimated from Equation (8). Using normal theory, we find the conditional distribution:

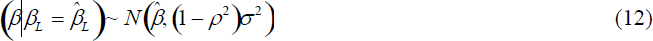

The distribution (12) is of interest, since, given the mean and variance of 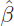, we can evaluate a new *t*-statistic

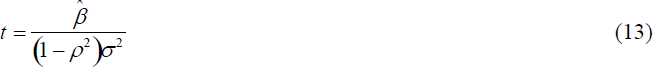

In order to do so, we will first estimate the expression (1−*ρ*^2^) from the distribution of 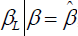, by assuming that

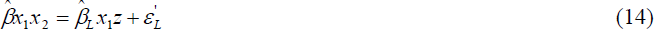

and by performing the latent variable regression (Equations (6)) on Equation (14) and then estimating the standard error using Equation (8). The assumption of Equation (14) is that only the genetic variance of the phenotype is explained by the latent interaction. With the new *t*-statistic in Equation (13) we are able to compute the *p*-value for the interaction of two SNPs, given that one of the SNPs had a significant interaction with a latent partner in the first stage of the procedure. We finally apply Bonferroni correction for the number of tests performed [51].

### Receiver operating characteristic (ROC) analysis

With the ROC analysis we are evaluating the performance of feature (SNP) classification. By fixing a threshold on the *p*-value, each method will classify the SNPs as either selected or not. The positive set consists of the two SNPs actually generating the phenotype. True positive and false positive rates were computed at a range of significance thresholds (*t*-test *p*-values) between zero and one. The area under the curve was computed using trapezoidal approximations. The analysis was performed in R.

### Dataset simulation

For genetic simulations, we used GenomeSIMLA [52]. We first simulated a population of genotypes based on human chromosome 22 as follows. For the 6,031 chromosome-22 SNPs found on the Affymetrix SNP Array 500K, we initialized a halplotype pool using the MAF values available on GenomeSIMLA’s website (http://ritchielab.psu.edu/ritchielab/software/genomesimla-downloads/). The initial population underwent random mating, genetic drift and recombination across 1,000 subsequent generations. The recombination frequencies were estimated by mapping the physical distances between the SNPs to genetic distances. The population growth was controlled using a logistic growth model. While the population is small, LD will develop and as the population grows larger the LD will diminish [52]. We sampled our datasets at two generations. For the comparison of LIMEpi to the exhaustive and marginal methods we sampled 100 datasets at generation 750. To study the effect of the overall LD in the sample on the second-stage time complexity, we sampled another 100 datasets at generation 500, where the population size was much smaller. To study the second-stage runtime as a function of the dimensionality of the dataset, we performed another independent simulation based on the entire Affymetrix SNP Array 500K. Within the limits of our computational resources, we grew a smaller population and sampled ten datasets at generation 750. The datasets from the above simulations underwent quality control, where we removed fixed alleles and alleles that switched from minor to major across generations. This filtering resulted in 3,395 SNPs for the chromosome 22 datasets and 261,858 SNPs for the whole-genome datasets. For each dataset above, we sampled 2,000 individuals from the pool. Phenotypes were simulated with various model parameter settings, but with fixed interaction coefficient *β* = 1.

### Working memory experiments and data

A total of 1,947 healthy young subjects (1,243 females, 704 males; mean age: 22.48 y; median age, 22 y; range, 18–35 y) were included in the study. The ethics committee of the Canton of Basel approved the experiments. Written informed consent was obtained from all subjects before participation.

## Authors’ contributions

JZK developed the statistical model, carried out the data analysis and drafted the manuscript. CV carried out the biological experiments and preliminary data analysis, helped to interpret biological results and helped to draft the manuscript. AH carried out the biological experiments, helped to interpret biological results and helped to draft the manuscript. DdQ conceived the study and participated in the study design and coordination and helped to draft the manuscript. AP conceived the study and participated in its design and coordination and helped to draft the manuscript. NB conceived the study and participated in its design and coordination, developed the statistical model and helped to draft the manuscript. All authors read and approved the final manuscript.

## Acknowledgements

This work was supported by the Swiss National Science Foundation CRSI33_130080.

